# Mutation in Abl kinase with altered drug binding kinetics indicates a novel mechanism of imatinib resistance

**DOI:** 10.1101/2021.06.28.449968

**Authors:** Agatha Lyczek, Benedict Tilman Berger, Aziz M. Rangwala, YiTing Paung, Jessica Tom, Hannah Philipose, Jiaye Guo, Steven K. Albanese, Matthew B. Robers, Stefan Knapp, John D. Chodera, Markus A. Seeliger

## Abstract

Protein kinase inhibitors are potent anti-cancer therapeutics (1). For example, the Bcr-Abl kinase inhibitor imatinib decreases mortality for Chronic Myeloid Leukemia (CML) by 80% (2, 3), but 22-41% of patients acquire resistance to imatinib (4). About 70% of relapsed patients harbor mutations in the Bcr-Abl kinase domain (5), in which more than a hundred different mutations have been identified (6–8). Some mutations are located near the imatinib binding site and cause resistance through altered interactions with the drug. However, many resistance mutations are located far from the drug binding site (9) and it remains unclear how these mutations confer resistance. Additionally, earlier studies on small sets of patient-derived imatinib resistance mutations indicated that some of these mutant proteins were in fact sensitive to imatinib in cellular and biochemical studies (10). Here, we surveyed the resistance of 94 patient-derived Abl kinase domain mutations annotated as disease-relevant or resistance-causing using an engagement assay in live cells. We found that only two-thirds of mutations weaken imatinib affinity by more than two-fold compared to Abl wild type. Surprisingly, one-third of mutations in Abl kinase domain still remain sensitive to imatinib and bind with similar or higher affinity than wild type. Intriguingly, we identified a clinical Abl mutation that binds imatinib with wild type-like affinity but dissociates from imatinib three times faster. Given the relevance of residence time for drug efficacy (11–14), mutations that alter binding kinetics could cause resistance in the non-equilibrium environment of the body where drug export and clearance play critical roles.

**Significance:** We performed the first in cell screen of imatinib binding against a library of Abl kinase mutants derived from patients with imatinib-resistant CML. The majority of mutations readily bind imatinib, posing the question of how these mutations cause resistance in patients. We identified a kinetic mutant that binds imatinib with wild type affinity but dissociates considerably faster from the mutant kinase. Using NMR and molecular dynamics, we found that this mutation increases the conformational dynamics of the mutant protein, linking conformational dynamics of the protein to drug dissociation. The results underline the importance of drug dissociation kinetics for drug efficacy and propose a novel kinetic resistance mechanism that may be targetable by altering drug treatment schedules.

## Introduction

Imatinib, a selective inhibitor of Bcr-Abl kinase, reduced the mortality from Chronic Myelogenous Leukemia (CML) by 80% within a decade of its FDA approval in 2001 (15, 16). Of the patients treated with imatinib in chronic phase, 95% achieve a complete hematologic remission and 60% a major cytogenetic response. However, most patients in blast crisis either fail to respond or quickly relapse following an initial response to imatinib. Mutations within the Bcr-Abl kinase domain are the most common cause for relapse (17). Over a thousand substitution mutations throughout the kinase domain have been identified in relapsed patients and are presumed to confer resistance to imatinib therapy (9, 18). Most of these mutations are located in areas of the kinase domain that are involved in imatinib binding, including mutations at the imatinib binding pocket, nucleotide-binding (P-loop) mutations, and activation loop (A-loop) mutations (19). Interestingly, more than 50% of the mutations in the COSMIC (20) or OncoKB (21) databases lie outside these areas. Early studies on small sets of kinase domain mutants showed that some clinically observed imatinib resistance mutations lost sensitivity to imatinib, while others showed little or no change (10). Furthermore, for some of these mutants, the biochemical and cellular imatinib sensitivities do not correlate. For example, the Abl E355G mutant remains sensitive to imatinib in biochemical assays, but only manifests its resistance in the context of cells (10). This observation could be explained based on several factors: (i) the purified mutant kinase constructs lack domains that facilitate resistance in the context of full length Bcr-Abl expressed in cells, (ii) mutations may increase kinase activity or biological stability in cells, or (iii) mutations could increase cellular autophosphorylation of Bcr-Abl (22).

Here, we propose a novel and general mechanism that mutations can cause resistance not by altering the equilibrium binding affinity, but rather by altering binding and dissociation kinetics. To illuminate this mechanism, we characterize a clinically observed mutation with significantly accelerated dissociation kinetics associated with relapse from imatinib therapy.

In contrast to most biochemical and cellular assays that employ a constant drug concentration over time (9, 10, 18), the plasma drug concentration of patients fluctuates over time due to drug pharmacodynamics and pharmacokinetics (11, 23). Under constant drug concentration, the affinity of the drug for the receptor is the major contributor to drug efficacy; however, in the non-equilibrium environment of the body, drug binding and dissociation are coupled to other dynamic processes, such as drug export and excretion that can limit drug efficacy. Drug-target residence time, defined as inverse of the dissociation rate, has emerged as an excellent predictor of drug efficacy in certain contexts (13, 24). Strategies aimed at prolonging drug-target residence time and minimizing off-target binding (i.e. kinetic selectivity) can drive drug efficacy (13, 24). Indeed, many successful kinase inhibitors such as imatinib and lapatinib have long residence times during which they elicit a pharmacological effect (11, 12, 25–27). An earlier study on Bruton’s tyrosine kinase (BTK) inhibitors showed that faster inhibitor dissociation correlated with lower efficacy (28). Based on these findings, we propose here that mutations can cause drug resistance by increasing dissociation rates, thereby decreasing drug residence times and reducing target inhibition in the non-equilibrium environment of the cell.

Here, we test how 94 patient-derived and resistance associated Abl mutants affect imatinib affinity and binding kinetics in live cells. We use a cell-based assay technique (NanoBRET^TM^) that employs bioluminescence resonance energy transfer (BRET) to directly measure molecular interactions between inhibitors and their kinase target in real-time (29, 30). This workflow represents the first broad-spectrum analysis of the target engagement characteristics of the full range of mutations in a clinically relevant kinase. We envision that this will serve as a paradigm for further investigation of analogous mutations in other kinases such as EGFR.

This technique has been recently used to study libraries of inhibitors against 178 full-length kinases (31). We present the first study in which a library of resistance-associated mutant kinases is screened for target engagement and residence time in live cells. We find that two-thirds of the tested clinically observed imatinib resistance mutations in Abl kinase significantly reduce drug affinity, whereas one-third bind imatinib with similar or tighter affinity than Abl wild type. We identified an imatinib resistance mutation, Abl N368S that has similar imatinib affinity as Abl wt, but increased imatinib dissociation rates. We propose here that clinical Abl mutations may cause drug resistance by suppressing the protracted residence time of imatinib in a pathophysiological setting.

## Results

To measure kinase engagement in live cells, we used a recently described technique that employs bioluminescence resonance energy transfer (NanoBRET) as a proximity-based measure of imatinib binding (32). Previously, this technique has successfully been applied to quantify compound engagement over a number of intracellular target proteins (29, 30). Briefly, a reporter complex forms inside live cells when a cell-permeable fluorescent probe molecule (tracer) binds to the ATP-binding site of a luciferase-tagged full-length target protein. In the presence of a cell-permeable luciferase substrate, the reporter complex emits a BRET signal. The addition of an unlabeled test compound (e.g. imatinib or dasatinib) disrupts the reporter complex and decreases the BRET signal. Our NanoBRET tracers were based on ATP-competitive kinase inhibitors that directly compete with the tested inhibitors for the same binding site on the kinase (31).

### Two-thirds of imatinib resistance-associated mutations bind imatinib with weaker affinity than Abl wild-type

We selected 94 mutations in Abl kinase domain mutations from the COSMIC (20) or OncoKB (21) databases that were annotated as disease relevant (OncoKB) or resistance causing (COSMIC) (**Table S1**). Some mutations are located in the ATP-binding site where they are likely to interfere with imatinib binding; however, many mutations are distal to the imatinib-binding site and their resistance mechanism is unclear (**Figure 1A**). We introduced individual point mutations to full-length Abl kinase fused to the C-terminus of NanoLuc luciferase (Abl1-Nluc) for expression in HEK293T cells.

**Figure 1.**
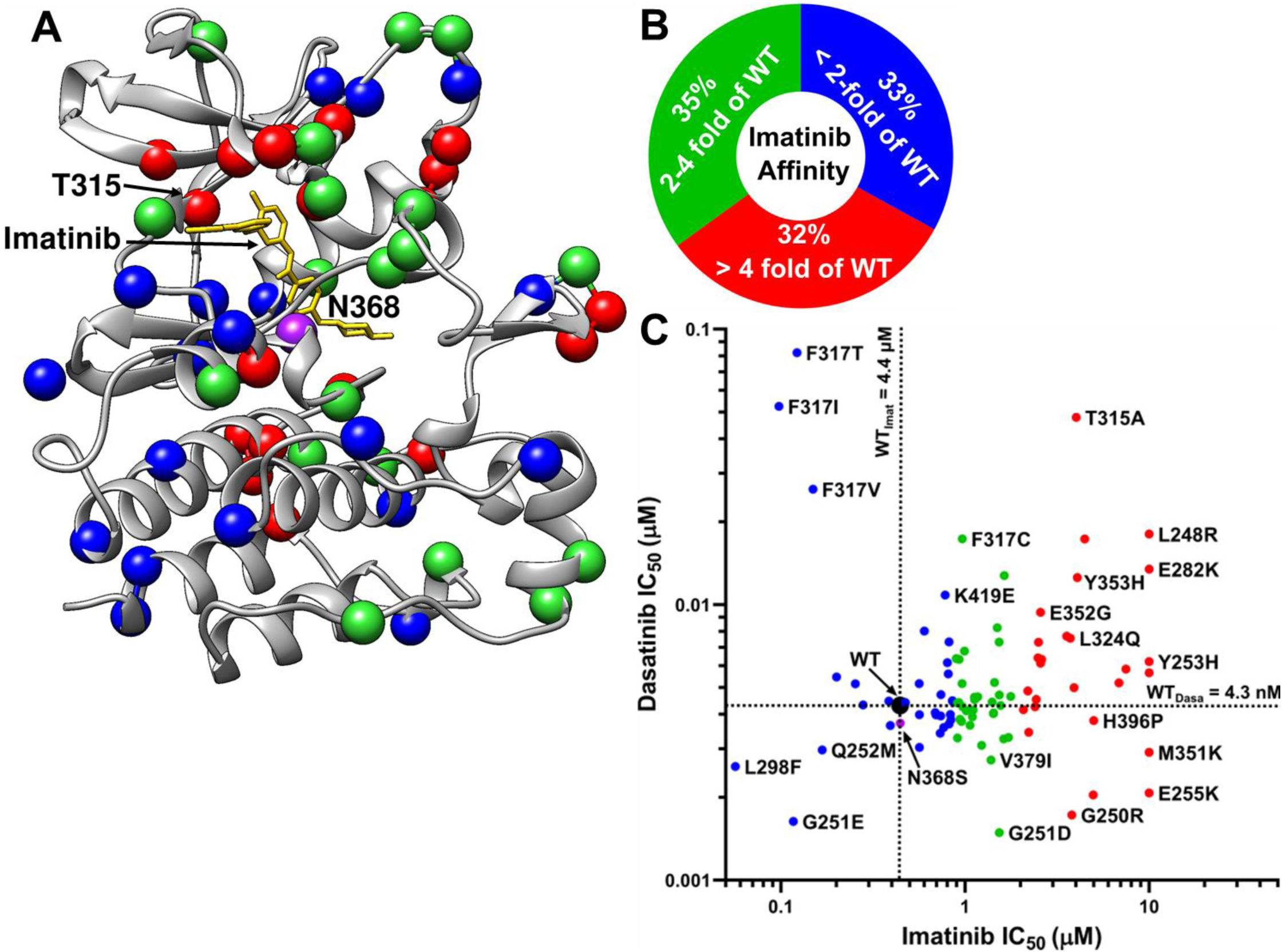
In-cell inhibitor binding affinities of mutant panel. (A) Structure of the Abl kinase domain (PDB-entry: 2HYY). The locations of 94 patient-derived Abl mutants associated with resistance are indicated as spheres. The color of the spheres indicates the level of imatinib resistance conferred by the mutation: blue – affinity of mutant is less than twofold weaker than WT, green – affinity is between two- and four-fold weaker than WT, red – affinity is more than eight-fold weaker than WT. (B) Fraction of mutations in different resistance classes. The colors correspond to panel A. (C) Imatinib and dasatinib resistance distribution of Abl mutants determined by in-cell NanoBRET target engagement assay. Colors correspond to panel A. Confidence of values is ± 23% based off of wt standard deviation, n=2.

Imatinib bound to Abl wt with a cellular affinity (IC_50_) of 0.44 ± 0.10 μM at equilibrium. For two-thirds of resistance-associated mutants (63 out of 94), the affinity for imatinib was weakened at least two-fold (IC_50_ ≥ 0.8 μM) (**Figure 1B&C**). At the standard dose of 400 mg daily, imatinib’s free serum concentration fluctuates from a mean maximum concentration of 4.4 ± 1.3 μM to a mean plasma trough concentration of 2 ± 1.4 μM (33). This indicates that in a population of patients, the trough plasma concentration would fall as low as 0.6 μM, which is below the IC_50_ of one-third of the tested mutants. The mutants that most drastically compromised imatinib affinity included the gatekeeper mutant and P-loop mutations, which had more than 8-fold higher imatinib IC_50_ values (**Table S2**). We were surprised to find that one-third of imatinib resistance mutations bound imatinib equallyor more potently than wt. The mutation that most drastically *increased* imatinib affinity was Abl L298F, which increased imatinib affinity four-fold over Abl wt (IC_50_ = 0.1 μM).

About 85% of mutants (80 out of 94) bound to dasatinib with similar affinity as Abl wt (cellular IC_50_ of 0.0043 μM) (**Figure 1B**). Only 15% (14 out of 94) of all the mutations tested had two-fold weaker dasatinib affinity than Abl wt, consistent with the clinical use of dasatinib as a second line therapeutic following emergence of imatinib resistance. Furthermore, mutations with weaker dasatinib affinity also showed weaker imatinib affinity. Exceptions to this were the mutations at F317, with F317I/T/V all remaining sensitive to imatinib, despite compromising dasatinib affinity (**Figure 1C**).

### Abl N368S binds imatinib with high affinity but dissociates more rapidly

Since a large fraction (31 out of 94) of imatinib resistance mutations still bound imatinib with affinity similar to wild type Abl, we determined the effect of mutation on cellular imatinib dissociation kinetics. Our rationale was that mutations might cause resistance by altering drug affinity or dissociation kinetics. We used NanoBRET assays to measure the real-time imatinib and dasatinib dissociation kinetics of mutants with wt-like imatinib affinities in live cells.

(**Figure 2 A&B**). In brief, Abl kinase was expressed in HEK293T cells, pre-loaded with imatinib or dasatinib and excess drug was washed out. An excess of fluorescent tracer was added that bound to Abl as imatinib/dasatinib dissociated resulting in an increased BRET signal.

**Figure 2.**
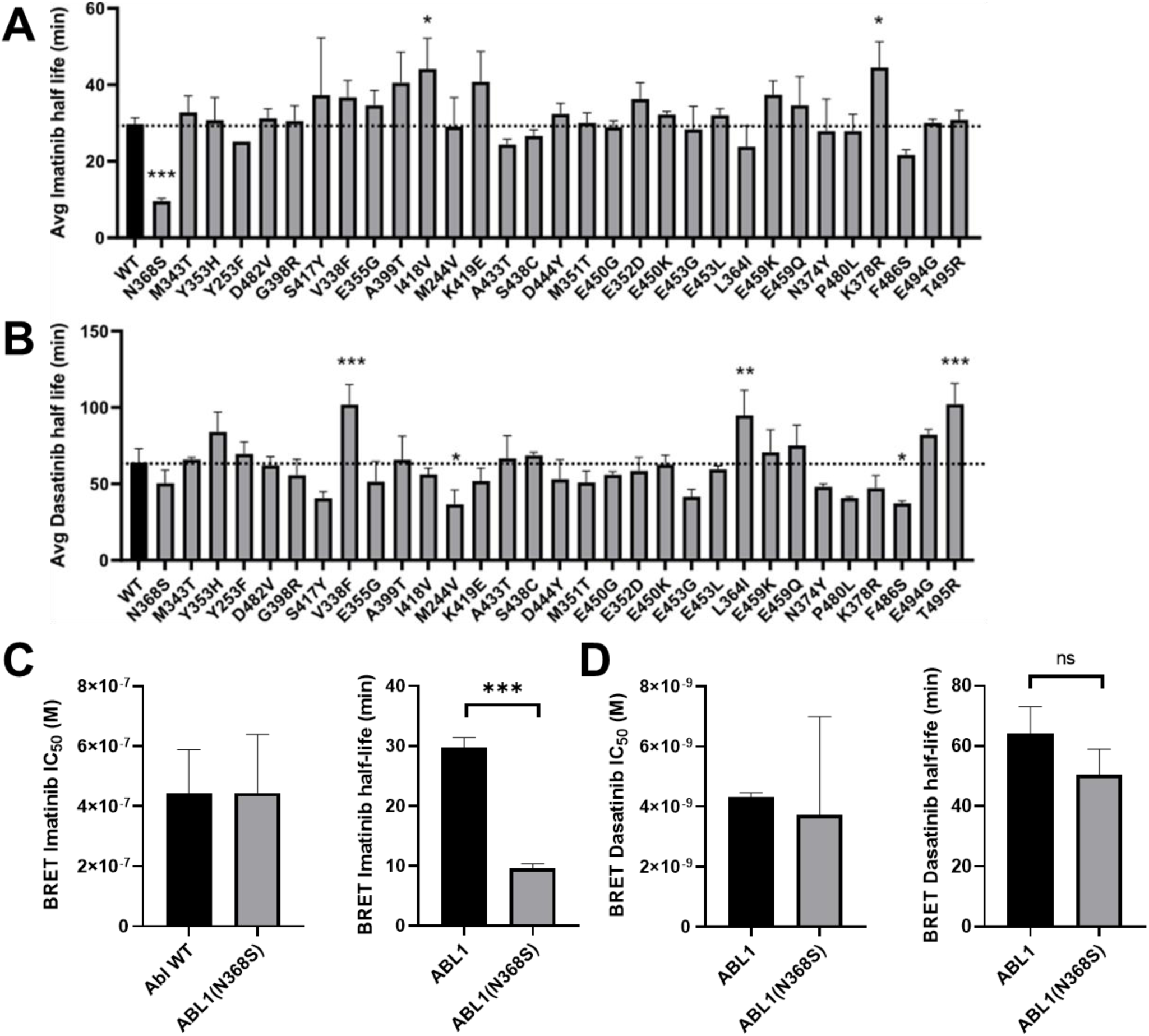
In-cell dissociation kinetics. (A) Abl mutants were characterized for imatinib dissociation kinetics using NanoBRET assay in live cells. (B) Dissociation kinetics of dasatinib from Abl mutants. The novel Abl N368S mutant was characterized for (C) imatinib and (D) dasatinib binding affinities and dissociation kinetics by NanoBRET in live cells. * p < 0.05, ** p < 0.01, *** p < 0.001

Imatinib and dasatinib dissociated slowly from Abl wt with half-lives of 29.7 ± 1.7 min and 64.1 ± 8.9 min, respectively. We found that the dissociation kinetics of dasatinib and imatinib were very similar among mutant proteins, and most mutations changed the dissociation rates constant relative to WT by less than 40 % (0.67 – 1.38 fold for imatinib and 0.63 – 1.75 fold for dasatinib).

Only one mutation, Abl N368S, a mutation that has only been reported once (34), accelerated imatinib dissociation kinetics by more than two-fold. Abl N368S bound imatinib with similar affinity to wt (0.44 ± 0.2 μM, 0.44 ± 0.14 μM respectively), but the imatinib dissociation rate was three times faster (half-life 9.6 ± 0.7 min) than wt (**Figure 2C**). This effect was specific to imatinib, as dasatinib dissociated from Abl N368S and Abl wt with similar kinetics (50.5 ± 8.4 min and 64.1 ± 8.9 min, respectively) (**Figure 2D**).

### Enzymatic activity of N368S kinase domain requires activation loop phosphorylation

The asparagine at position 368 in Abl WT is nearly universally conserved among kinases (**Figure 3A**) (35). The side chain of N368 forms hydrogen bonds with D381 and the catalytic aspartate D363 when the kinase is in the DFG-Asp-in conformation (PDB-entry 2G2I) (**Figure 3B**). Both D381 and D363 coordinate Mg^2+^/ATP and they are required for catalysis (36, 37). Therefore, we speculated that N368 could be critical for enzyme activity as well.

**Figure 3.**
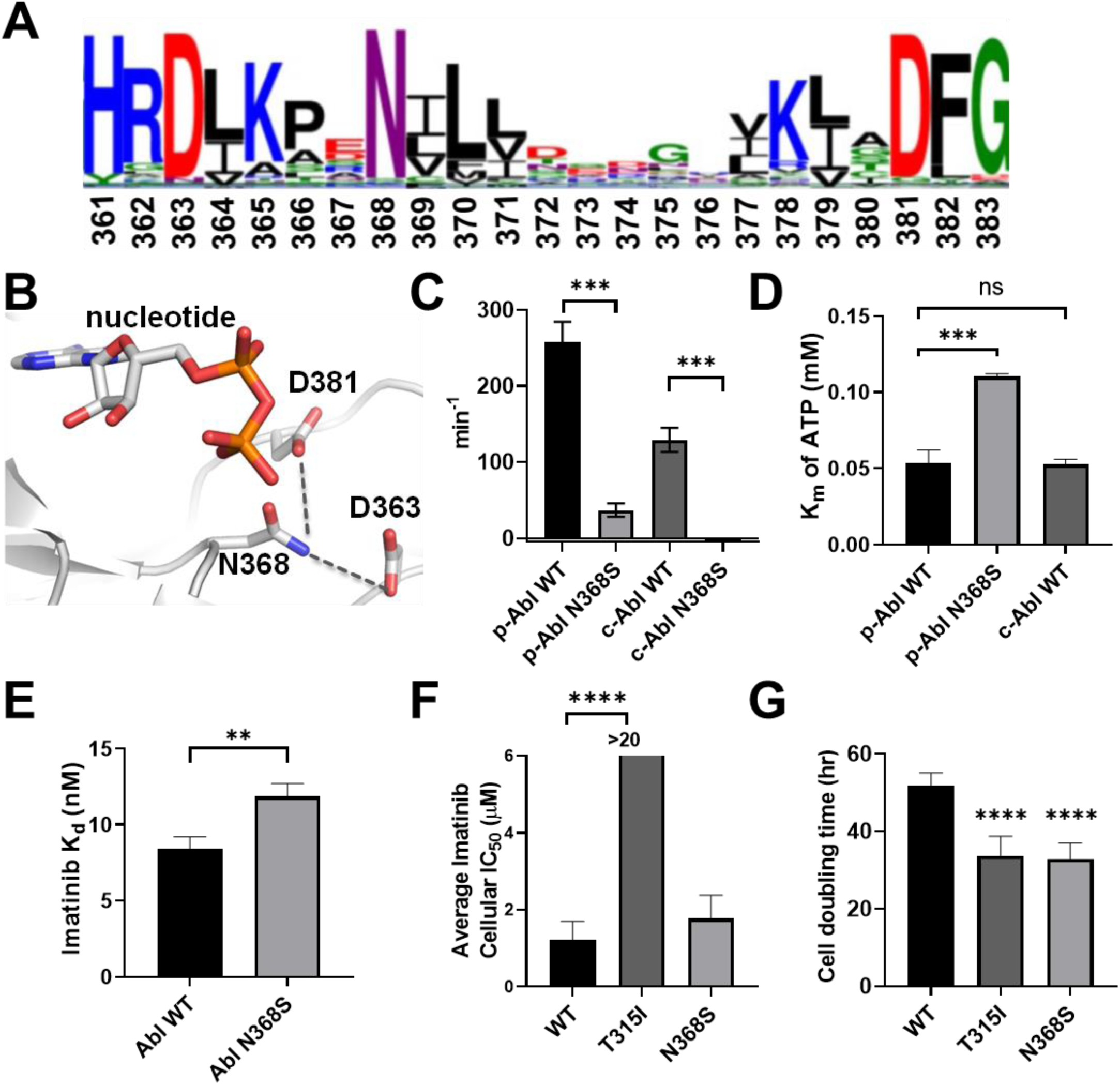
Characterization of the Abl N368S mutation. (A) Sequence alignment of human protein kinases represented as weblogo. The size of the letters reflects the degree of conservation and numbering is according to Abl1A numbering. Asparagine 368 is as highly conserved as the catalytic HRD and the DFG motif. (B) Structural overview of Abl N368 in the active conformation (PDB-entry 2G2I). The sidechain of N368 is near the nucleotide and is positioned to form hydrogen bonds with the catalytic aspartate D363 and D381 in the DFG-motif. (C) Kinase activity of purified Abl N368S is undetectable in the absence of activation loop phosphorylation (designated as pAbl). (D) The Michaelis-Menten constant of purified pAbl N368S for ATP is two-fold higher than for pAbl WT. (E) The affinity of purified of Abl N368S for imatinib determined by a spectroscopic binding assay is slightly weaker than for Abl WT. (E,F) BaF3 cells expressing BCR-Abl WT, T315I or N368S. BaF3 cells expressing N368S are equally sensitive to imatinib as cells expressing BCR-Abl WT (E). (F) BaF3 cells expressing imatinib resistance mutations T315I or N368S exhibit a faster doubling time in the absence of imatinib. * p < 0.05, ** p < 0.01, *** p < 0.001, **** p < 0.0001

We expressed and purified unphosphorylated Abl kinase domain (KD) to characterize the N368S mutant biochemically. Abl N368S was predicted to be stable according to FoldX calculations and the experimental melting temperature of Abl N368S (46.05 ± 0.13 °C) was virtually identical to that of Abl wt (45.69± 0.07 °C) (**Figure S1**). Next, we determined how the N368S mutation affected the affinity of Abl kinase domain for imatinib. The intrinsic protein fluorescence of Abl decreased upon imatinib binding and therefore can be used as a readout for imatinib binding (38). Consistent with our cellular NanoBRET assays, we found that the dissociation constant (K_d_) of imatinib for purified Abl N386S (K_d_ = 11.8 ± 0.9 nM) was only 1.5-fold higher than that of Abl wt (8.4 ± 0.8 nM), making it unlikely that this difference contributes to inhibitor efficacy *in vivo* (**Figure 3E**).

When evaluating the enzymatic activity of Abl N368S kinase domain we were surprised that unphosphorylated Abl N368S had no detectable catalytic activity (k_cat_ unmeasurable) compared to Abl wt (k_cat_ = 129.0 ± 31.5 min^-1^). However, activation loop phosphorylation of N368S by Hck increased kinase activity (k_cat_ = 36.9 ± 17.6 min^-1^). This phosphorylation activity was still 7-fold lower than the activity of Abl wt phosphorylated at the activation loop by Hck (k_cat_ 258.0 ± 52.2 min^-1^) (**Figure 3C**). Next, we determined how the N368S mutation affected the Michaelis-Menten constant (K_m_) of Abl kinase domain for ATP. We found that activation loop phosphorylated Abl (pAbl) N368S (K_m_^ATP^ = 110.3 ± 1.9 μM) had greater than two-fold K_m_ for ATP as pAbl wt and non-phosphorylated Abl wt (K_m_^ATP^ = 53.2 ± 8.9 μM,) (**Figure 3D&F**). This suggested that the mutation had a two-fold lower ATP affinity and therefore increased competition between ATP and imatinib was not the mechanism behind N368S’s resistance phenotype.

### N368S affects drug dissociation kinetics of isolated kinase domain

It is conceivable that regulatory domains, interactions with cellular proteins or ligands, or biological processes such as protein turnover facilitate the faster dissociation of imatinib from Abl N368S in cells. Thus, we determined whether purified Abl N368S kinase domain also showed an increased rate of imatinib dissociation using a competition assay between the two ATP-competitive drugs bosutinib and imatinib. Upon binding to Abl kinase bosutinib fluorescence at 450 nm increases, making it a useful probe to monitor inhibitor binding to kinases (39, 40). Briefly, after formation of Abl-imatinib complexes, we add a five-fold molar excess of bosutinib. Bosutinib competes with imatinib for the ATP-binding site, and upon binding, there is an increase in fluorescence at 450 nm. We found that imatinib dissociated two-fold faster from Abl N368S than from Abl wt at 25°C, 32°C, and 37°C (**Figure 4A**). Similarly, imatinib dissociated two-fold faster from phosphorylated Abl N368S than from Abl wt at 25, 32, and 37°C (**Figure 4B**). These results suggested that faster imatinib dissociation from full-length Abl N368S in cells is directly caused by an increased inhibitor off-rate from the kinase domain.

**Figure 4.**
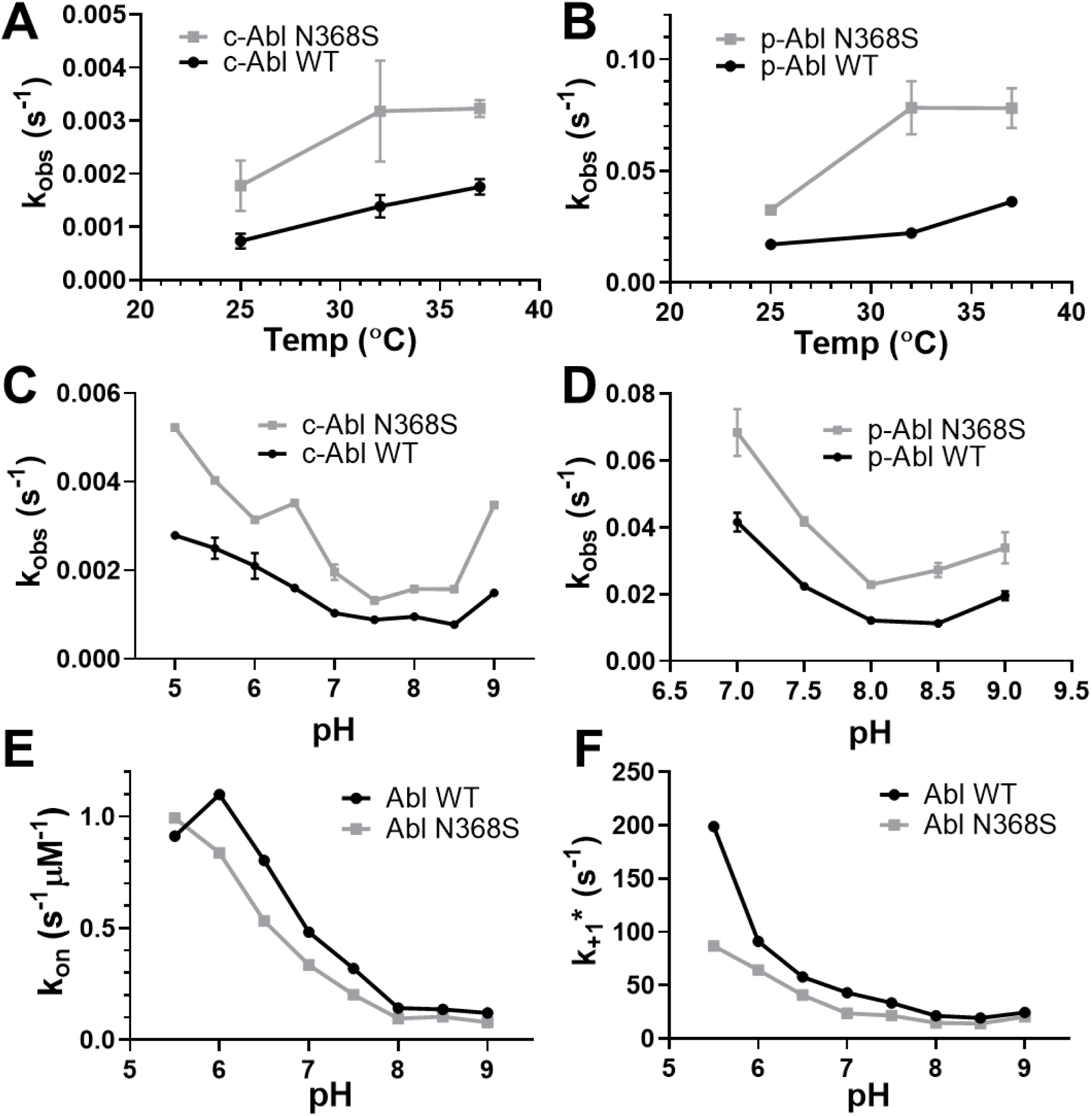
In vitro kinetics of imatinib binding and dissociation from purified Abl kinase domain. (A, B) Dissociation kinetics of imatinib from unphosphorylated (A) and activation loop phosphorylated (B) Abl WT and N368S at various temperatures. (C, D) Dissociation of imatinib from unphosphorylated (C) and activation loop phosphorylated (D) Abl WT and N368S at various pH values. (E, F) Binding kinetics of imatinib to Abl WT and N368S at various pH values yielding the rate constant of imatinib binding (E) and the rate constant of the DFG-flip (F).

### Faster imatinib dissociation from Abl N368S is pH independent

We showed previously that the rate of imatinib binding to Abl kinase domain is pH dependent. For imatinib to bind, the side chain of Asp381 in the DFG motif must rotate from a position where it faces into the active site (DFG-Asp-in) to an orientation where it faces the solvent (DFG-Asp-out); a process referred to as the DFG-flip. Protonation of Asp381 reduces the energetic barrier for the 180° crankshaft-like rotation of the DFG motif and increases the imatinib binding rate without changing the affinity (41). This indicates that the DFG-flip may be the rate-limiting step for the imatinib binding process.

Next, we wanted to investigate whether the N368S mutation affects the DFG-flip and whether the DFG-flip remained the rate limiting step for imatinib dissociation. We determined how pH affected the kinetics of imatinib binding to Abl wt and N368S using our previously described fast transient stopped-flow assay (38). We found that the rate constants for imatinib binding (*k*_on_) and the DFG-flip (*k*_+1_*) were about 30% slower for N368S than for Abl wt (*k*_on_ = 0.143 ± 0.076 μM^-1^s^-1^ and *k*_+1_* = 21.4 s^-1^ respectively at pH 8.0) and Abl N368S (*k*_on_ = 0.096 ± 0.006 μM^-1^s^-1^ and *k*_+1_* = 14.6 s^-1^ respectively at pH 8.0) across the pH range (**Figure 4E, F**) (41).

Next, we tested whether the dissociation rate of imatinib from Abl is pH dependent, which would indicate that the DFG-flip was rate limiting for imatinib dissociation. We measured imatinib dissociation kinetics at different pH conditions ranging from pH 5.0 to 9.0. We found that for both Abl wt and N368S, imatinib dissociation kinetics were independent of pH. This makes it unlikely that the protonation-sensitive DFG-flip was a rate limiting step in imatinib dissociation. Imatinib dissociated from Abl N368S two-fold faster than from Abl wt across the tested pH range (**Figure 4C**). Likewise, imatinib dissociated two-fold faster from phosphorylated Abl N368S than from wt across the tested pH range (**Figure 4D**).

Taken together, these data show that compared to Abl wt, faster imatinib dissociation from N368S is not facilitated by a faster DFG-flip, which in fact is slightly slower in N368S. Instead, the N368S mutation mainly alters the dissociation process.

### Ba/F3 cells expressing Bcr-Abl N368S proliferate and remain sensitive to imatinib under constant drug concentration

Next, we determined whether the Abl N368S mutation had any effect on cellular proliferation. Upon expression of active human Bcr-Abl, murine hematopoietic Ba/F3 cells proliferate independently of IL-3 (9). Ba/F3 cells expressing the Bcr-Abl1 N368S mutant proliferated in the absence of IL-3, which is further evidence that the Bcr-Abl N368S mutant is active in a cellular context. We determined the imatinib cellular IC_50_ for three cell lines expressing Bcr-Abl wt or resistance-associated mutations (wt, T315I, N368S). Imatinib inhibited the proliferation of cells expressing Bcr-Abl wt and N368S with IC_50_ values of 1.23 ± 0.47 μM and 1.78 ± 0.60 μM, respectively (**Figure 3F**). We used a cell line expressing the T315I gatekeeper mutation as a control and found that it was highly resistant to imatinib as expected (IC_50_ = 36.6 ± 12.0 μM). These results were in excellent agreement with the NanoBRET target engagement results for wt, N368S and T315I (IC_50_-values of 0.44 ± 0.14 μM, 0.44 ± 0.2 μM, >10 μM respectively), considering the use of different cell lines and Abl construct in the different assays.

We also determined whether the presence of imatinib resistance-associated mutations affects cell doubling time. We found that doubling times were statistically significantly shorter for Ba/F3 cells expressing Abl T315I (33.7 ± 2.0 hrs) and N368S (32.9 ± 1.7 hrs) than for cells expressing Abl wt (51.9 ± 1.1 hrs) (**Figure 3G**). Because of the exponential growth of cells, the 1.6-fold faster doubling times results in 10-times more cells expressing N368S than wt after 300 hrs. Taken together, Bcr-Abl N368S accelerates cell proliferation and is inhibited by imatinib under constant drug concentration similarly to wt.

### The H-bond interaction between N368 and D381 is only observed in the DFG-in conformation

Since drug binding and dissociation is a dynamic process, we speculated that protein dynamics could be affected by the N368S mutation. Therefore, we performed twelve 500 ns MD simulations with a self-adjusted mixture sampling (SAMS) enhanced sampling scheme (42) that accelerates sampling of DFG-flip events. We chose two different starting structures: Abl in the DFG-out conformation, bound to imatinib (PDB entry: 2HYY), and Abl in the DFG-in conformation bound to dasatinib (PDB entry 2GQG). Simulations were run for each of these starting models in the presence and absence of the corresponding ligand, and with and without the N368S mutation. We employed two slightly different SAMS biasing strategies which we describe in the *Materials and Methods* section.

Among the twelve simulations, we observed five DFG flips in the apo simulations and none in the presence of ligands (**Table S3**). In four simulations of Abl wt and N368S, the aspartate of the DFG-motif flipped from a solvent-exposed, outward orientation to an orientation facing into the active site of the kinase (out-to-in flip). In one additional simulation of N368S, the DFG-motif flipped in the reverse direction (in-to-out). This indicated that N368S could perform the DFG flip in both directions.

Next, we investigated the H-bonding of Abl wt and N368S within the active site. We found that the side chain of N368 formed an H-bond with DFG-Asp only in the DFG-Asp in conformation (**Figure 5 A, B**), and those interactions were absent in the N368S mutant. The H-bond between N368 and D381 likely stabilized the DFG-in (active) conformation and acted as a kinetic barrier for the DFG flip. Conversely, loss of the H-bond in Abl N368S would be expected to destabilize the active conformation, consistent with the low specific activity of N368S. Next, we investigated how the N368S mutation affected the solution dynamics of the activation loop. Based on the hydrogen-bonding pattern of Asn368, we hypothesized that Asn368 could anchor the amino-terminal end of the activation loop and helix αC. We therefore compared ^1^H-^15^N NMR peak intensities for WT and N368S (**Figure 5C, Figure S3**). Sharper NMR peaks leading to higher signal intensities correspond to increased solution dynamics. We found that several residues within the activation loop of N368S showed increased dynamics compared to WT. In particular, G383 in the DFG-motif exhibited a five-fold stronger signal relative to WT, indicating a sharp increase in dynamics. Similarly, all residues in helix αC that face the DFG motif showed increased dynamics in N368S. This observation is consistent with the notion that in the N368S mutant activation loop and helix αC become more dynamic. In a separate study, we have performed molecular dynamics (MD) simulations of the dissociation process of imatinib from Abl WT and N368S (43). These simulations quantitatively reproduce the experimentally observed imatinib dissociation rates for WT and N368S. The MD studies indicated that the flexibility of the activation loop and helix αC are crucial for imatinib dissociation, consistent with our experimental observations.

**Figure 5.**
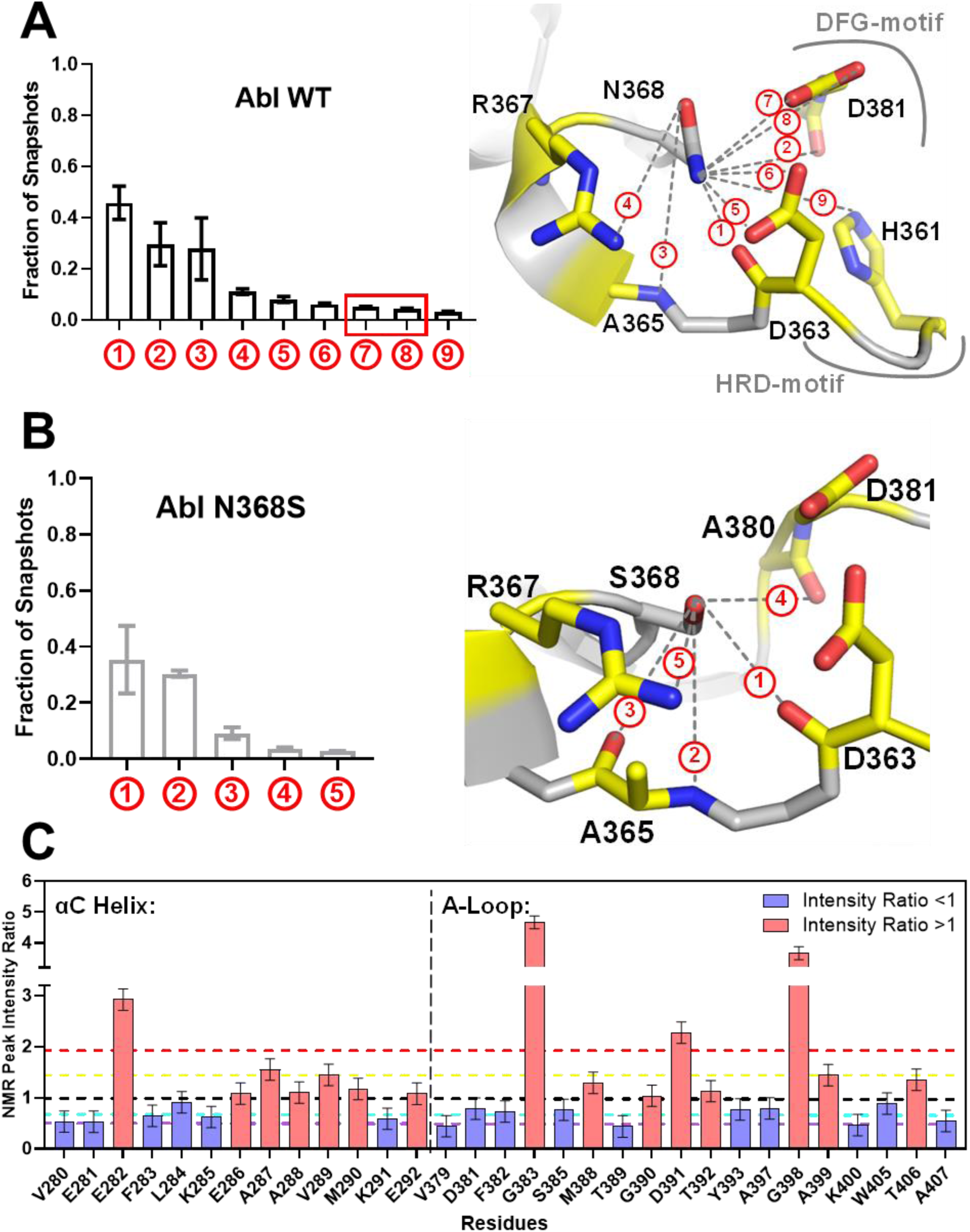
Molecular Dynamics and NMR analyses of the N368S mutant protein. (A) Fraction of snapshots from MD simulation in which the side chain of N368 engages in a hydrogen bond with neighboring atoms (left) illustrated in the structure of Abl kinase in the active conformation (PDB-entry: 2GQG) (right). (B) Fraction of snapshots from MD simulations in which the side chain of S368 engages with neighboring atoms (left) illustrated on the model of S368 in the structure of active Abl WT (PDB-entry: 2GQG). (C) NMR ^1^H-^15^N peak intensity ratios (I_N368S_/I_WT_) for residues in helix αC and the activation loop. Black line = ratio of 1, yellow line = mean of all values larger than 1, red line = 1 standard deviation from mean of values larger than 1, cyan line = mean of all values smaller than 1, purple line = 1 standard deviation from mean of values smaller than 1.

## Discussion

In this study we determined the target engagement of 94 resistance-associated mutations of Abl kinase for the cancer drugs imatinib and dasatinib in cells. Surprisingly, we found that one-third of imatinib resistance-associated mutations remain sensitive to imatinib and bind with similar affinity as Abl wt. One explanation could be that some mutations may not cause resistance to imatinib but they exist on the background of other resistance mechanisms such as upregulation of drug exporters, downregulation of drug importers, increased serum protein binding, or overexpression of Src kinase (44–46). Importantly, this would indicate that approximately one-third of mutations in the relevant databases are incorrectly assigned. If identified in patient samples, the presence of these mutations should not necessarily alter clinical standard of care. Imatinib could remain the first line therapy for patients harboring these mutations unless other clinical metrics indicate disease resistance to imatinib.

Potential limitations of this proof of principle pilot study include the use of full-length Abl instead of Bcr-Abl and the use of HEK293T cells instead of a more disease relevant hematopoietic cell line. Differences in the concentration, activity, and localization of other proteins and ligands in Bcr-Abl-positive leukemic cells could affect drug-target engagement. Therefore, we cannot exclude the possibility that some mutations may be more resistant to imatinib in the Bcr-Abl protein and in the context of leukemic cells. For example, mutations might sensitize Abl to tetramerization via the Bcr region or to activation loop phosphorylation through other kinases, leading to increased Abl activation loop phosphorylation and reduced imatinib affinity (47).

To explore novel molecular resistance mechanisms, we determined the cellular drug dissociation kinetics for 34 Abl mutants. Given that drug residence time is a better predictor for drug efficacy than affinity alone (11–13), we predicted that, conversely, mutations could reduce drug efficacy by reducing drug residence time. We found that Abl N368S had similar imatinib affinity compared to wt, but its imatinib dissociation was three times faster than wt. This observation posed the possibility of kinetic drug resistance caused by mutations. Such kinetic mutations may not be detectable in assays with constant drug concentration and their resistance phenotype may vary from patient to patient by other pharmacodynamic and pharmacokinetic parameters. Importantly, kinetic resistance mechanisms may be addressable by altering the treatment schedule, for example from a single daily dose to multiple doses.

Our biochemical and biophysical studies indicate that imatinib dissociation is not a simple reversal of the binding process. Consistent with such a mechanism, a recent computational study using adaptive sampling between conformational “milestones” proposes a dissociation pathway of imatinib from Abl where the kinase lingers in the DFG-out conformation via a temporary deformation of the N-lobe (48). To address the mechanism of N368S, we performed extensive simulations of imatinib dissociation from wild type and N368S kinase using infrequent metadynamics (43, 49). This method captured qualitatively and quantitatively the off-rates for both systems and demonstrated how the faster off-rate for the mutant is a direct consequence of enhanced protein flexibility in the DFG region.

To our knowledge, the work presented here is the first study to systematically evaluate a large panel of clinical mutations for drug engagement and changes in residence time in live cells. This workflow provides a template for broader evaluations of disease relevant mutations and how such mutations impact drug affinity and binding and dissociation kinetics.

With more clinical sequencing efforts of patient samples, we expect that databases on disease-related mutations will grow rapidly in size. Thus, our results indicate that the experimental characterization of mutant phenotypes in cells remains essential to distinguish mutations that are causal for observed clinical phenotypes versus those observed in the background of other biological processes.

## Materials and methods

### Protein Expression and Purification

The kinase domain of human c-Abl (UniProtID: P00519, residues 248– 531), human Abl 1a numbering (50) was cloned into pET28, modified to yield an N-terminal, TEV-cleavable His-tag (51). The kinase domain c-Abl mutations were introduced into the human c-Abl kinase domain by site-directed mutagenesis using the QuikChange II kit (Agilent) and verified by DNA sequencing.

Kinases were co-expressed in *Escherichia coli* BL21DE3 cells with full-length YopH phosphatase from *Yersinia pseudotuberculosis* (52) and GroEL/Trigger factor as previously described (38, 51). *E. coli* cells were grown in 2xYT media (GIBCO) at 37°C for 4 hours to a cell density of OD_600_=0.6. The temperature was subsequently reduced to 16°C and cells were induced with 1 mM IPTG and grown overnight. For generation of ^15^N-labeled protein for NMR studies, protein was expressed in M9 minimal media with ^15^N-NH_4_Cl (Cambridge Isotope labs) as the sole nitrogen source instead of 2xYT media. Kinases were purified by a Ni affinity column (HisTrap FF, GE Lifescience). The his-tag was cleaved by TEV protease. Kinases were then purified using anion exchange chromatography (HiTrap Q FF, GE Lifescience), and size-exclusion chromatography (S75, GE Lifescience). Proteins in 20 mM Tris (pH 8.0), 250 mM NaCl, 5% glycerol, 1 mM DTT were concentrated, frozen in liquid nitrogen, and stored at −80°C. Protein purity was at least 95% as determined by SDS-PAGE and Coomassie staining.

### NMR spectroscopy

Purified protein was concentrated to a final concentration of 300 µM by ultrafiltration and buffer exchanged into 50 mM Na/K pH 6.5, 50 mM NaCl, 5 mM DTT at 4000 rpm, 4 °C. Imatinib was added in five-fold excess during the buffer exchange. Deuterium oxide was added to 10% final concentration in the NMR samples for a lock signal. All backbone (^1^H)-^15^N heteronuclear NMR experiments were acquired at 27 °C on a Bruker Avance III HD spectrometer operating at a ^1^H frequency of 850 MHz equipped with a cryogenic probe. Standard Bruker pulse sequence for ^1^H-^15^N TROSY-HSQC, trosyf3gpphsi19.2, was used. Sample stability prior and post 2D ^1^H-^15^N NMR experiments was assessed by acquiring and comparing the relative intensity of the 1D ^1^H spectra prior and post experiments. The backbone assignments from BMRB were transferred to the Abl WT/N368S spectra (53). The intensity change was analyzed by NMRFam-Sparky (54), and the graphs were created using Prism GraphPad. The intensity ratios were calculated as the ratio of the respective cross peak height in N368S and WT. Differences in sample signal intensity between the Abl WT and N368S samples were calibrated as described previously (55).

### Tissue Culture

Human embryonic kidney 293 cells (HEK293T cells, ATCC) were grown in Dulbecco’s Modified Eagle media (DMEM, Thermofisher) with 10% Fetal Bovine Serum (FBS, Thermofisher) and 1% penicillin/streptomycin (Thermofisher). IL3-dependent murine pro-B cell (Ba/F3 cells, ATCC) were maintained in Roswell Park Memorial Institute (RPMI) media (Thermofisher) supplemented with 10% FBS and IL-3 and 1% penicillin/streptomycin. A murine B lymphocyte cell line, Wehi -231 cells (ATCC), was used to produce and harvest IL-3. Wehi cells were maintained in RPMI media supplemented with 10% FBS and 1% penicillin/streptomycin. The supernatant was harvested after Wehi cells reached a density of more that 2 million cells/mL confluent and media turned orange. IL-3 was harvested from the supernatant of Wehi cells and filtered using a 0.45 μm filter. Cells were kept in a humidified incubator at 37°C and 5% CO_2_. Media was changed every 3 days. Cells were split every 6 days with 0.25% trypsin (Thermofisher).

### Ba/F3 Cell line mutagenesis and cell line generation

Full-length p210 *Bcr-Abl* (BA) cDNA was sub-cloned into *pEYK3.1*, a retroviral vector to construct the *pEYKBA* plasmid (56). The *pEYKBA* plasmid was mutagenized by site-directed mutagenesis using the QuikChange II kit (Agilent) to create point mutations in the kinase domain of *Bcr-Abl*. The *pEYKBA* plasmids were transfected into 293T cells using FuGENE HD (Promega) as described in section 2.2.3. At 24 hours after transfection retroviral supernatants were recovered as previously described (9, 56). Interleukin 3-dependent hematopoietic pro-B cell line Ba/F3 (57) were transduced with the retroviral supernatant supplemented with 8 μg/ml of polybrene (Sigma) and IL-3. Ba/F3 were deprived of IL-3 two days post-infection to select for cells that had taken up the *pEYKBA* retrovirus.

### Ba/F3 Cell Viability assay to determine IC_50_

0.5 x 10^4^ Ba/F3 cells expressing mutant Bcr-Abl proteins were plated in each well of 96-well plates in RPMI medium containing 10% FBS and lacking IL-3 (total volume 100 μL). Imatinib was included in the media in increasing concentrations (final concentration ranging from 0 to 20 μM). Dasatinib was included in the media in increasing concentrations (final concentration ranging from 0 to 10 μM). Viable cell number was assessed 48 hr later using the WST-1 reagent (Roche), according to manufacturer’s specifications. Assays were performed in quadruplicate. A_450_ readings were plotted versus the logarithm of the concentration of drug and fit to a sigmoidal equation with the hill slope set to 1.0. The concentration of drug resulting in 50% maximal inhibition is reported as cellular IC_50_.

### NanoBRET Assays for target engagement

Bioluminescence resonance energy transfer (BRET) can be used as a proximity-based measure of drug binding in live cells (32). In brief, an energy transfer occurs between the 19-kDa NanoLuc luciferase (NanoLuc, Nluc)-tagged Abl target protein and a cell-permeable fluorescent energy transfer probe introduced to the cell culture medium. Imatinib binding results in competitive displacement of the probe and a loss of energy transfer. Thus, this technique provides a measure of intracellular drug-target engagement.

The NLuc-Abl1 full-length plasmid was a kind gift of Promega. The 94 Abl mutants were generated in the MacroLab at Berkeley (40). They used an existing library of Abl mutant constructs that were created using site directed mutagenesis. Using the original vector, they cloned in a dropout cassette in place of the sequence that contained the desired mutation sequences to generate an intermediate molecule. By digesting with AarI, the dropout cassette was removed. The mutant constructs were amplified with the appropriate primers to clone into the new plasmid backbone (again by digesting with AarI). Then they did Golden Gate cloning, a molecular cloning method that allows for simultaneous and directional assembly of multiple DNA fragments into a single piece using Type IIs restriction enzymes and T4 DNA ligase. AarI restriction enzyme was used to insert the PCR fragments into the new plasmid backbone. Plasmids were verified by sequencing.

For transfecting HEK293T cells, Nluc/Abl fusion constructs were mixed with carrier DNA (pGEM3ZF-, Promega) at a mass ratio of 1:10 (mass/ mass) to prepare a 10 μg/ml solution of DNA in OPTIMEM (Thermofisher). Subsequently, DNA was mixed with FuGENE HD (Promega) to form complexes according to the manufacturer’s protocol (Promega). DNA:FuGENE complexes were formed at a ratio of 1:3 (mg DNA per ml FuGENE). One part of the transfection complexes was then mixed with 20 parts (v/v) of HEK293T cells suspended at a density of 4×10^5^ in DMEM (Gibco). Cells were seeded into white, nonbinding polypropylene plates (Greiner 781207) and incubated for 20 h prior to experiments. All chemical inhibitors were prepared as concentrated stock solutions in dimethylsulphoxide (DMSO) (Selleckchem).

For generating tracer isotherms against targets expressed in cells, tracer compound was added to cells using a liquid dispenser, Echo 550 (Beckman) in a range of concentrations from 0 to 1 μM. Cells were then equilibrated with the tracer for two hours. After two hours, the 3X Complete Substrate and Inhibitor was added to cells. The 3X Complete Substrate and Inhibitor solution was prepared by mixing the NanoBRET Nano-Glo Substrate (Promega) and Extracellular NanoLuc Inhibitor (Promega) into OPTI-MEM. The 3X Complete Substrate and Inhibitor solution was then added to cells and luminescence was measured at 450 nm (donor emission) and 610 nm (acceptor emission). We identified a common tracer probe (K-4, Promega), suitable for all mutants, and determined its assay window (**Table S4**) and apparent affinity for all mutants (**Table S5**).

For determining imatinib and dasatinib isotherms, NanoBRET tracer was added to the cells at fixed concentrations (the tracer *IC*_50_ concentration for each respective Abl mutant). Cells were incubated with tracer and serially diluted test compound using a liquid dispenser, Echo 550 (Beckman) and incubated for 2 hours prior to read out to allow the binding to reach equilibrium. The 3X Complete Substrate and Inhibitor solution was then added to cells, and filtered luminescence was measured at 450 nm (donor emission) and 610 nm (acceptor emission). Background-corrected BRET ratios were determined by subtracting the BRET ratios of samples from the BRET ratios in the absence of tracer and test compound. BRET ratios were plotted as a function of the log(inhibitor concentration) to determine *IC*_50_ -the inhibitor concentration that provokes a response halfway between the maximal response and maximally inhibited response. To calculate *IC*_50_, equation (1) was used in GraphPad Prism 8, where the Hillslope describes the steepness of the curve is fixed to a value of -1.0, and the Top and Bottom describe the plateaus in the units of the Y axis (3-parameter curve fit).

### BRET Assays for residence time

For kinetic analysis of compound dissociation rates, HEK293T cells transfected with Nluc-Abl1 fusion constructs as described above were pre-equilibrated for 2 h with the respective 10x *IC*_50_ concentrations of test compound for each mutant (imatinib or dasatinib). Cells were subjected to a wash-out step where they were centrifuged, media was removed and fresh media was added to cells containing NanoBRET NanoGlo Substrate and tracer (at 20x the determined tracer Kd,app) diluting at least 100 times, kinetic BRET measurement was immediately performed over 2.5 hours at 30°C. BRET ratios as a function of time were fitted to a single exponential equation with or without a sloping baseline (**Figure S4**) to obtain the observed rate constant (*k*_obs_) and dissociation reaction half-life.

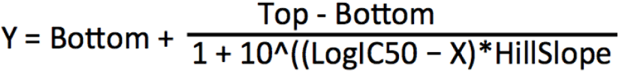

### Phosphorylation of kinase protein

Abl kinase domain was incubated at 4 °C overnight with a bacterially expressed construct of human Hck kinase containing the SH3-SH2 and kinase domains (0.2 mg/mL and 0.01 mg/mL respectively), ATP (2 mM), sodium orthovanadate (1 mM) 100 mM Tris (pH 8.0), and 10 mM MgCl_2_. Phosphorylation was confirmed by kinase activity assay.

### Kinase activity assay

Kinase activity was monitored using a continuous spectrophotometric assay as described before (58, 59). In brief, the consumption of ATP is coupled via the pyruvate kinase/lactate dehydrogenase enzyme reactions to the oxidation of NADH, which can be monitored through the decrease in absorption at 340 nm. The reactions were run under pseudo-steady state conditions as ATP was regenerated during the reaction to keep a steady concentration of ATP. Reactions contained 100 mM Tris (pH 8.0), 10 mM MgCl_2_, 0.5 mM ATP, 1 mM phosphoenolpyruvate, 0.21 mg/ mL NADH, 75 U/mL pyruvate kinase, 105 U/mL lactate dehydrogenase, and 0.5 mM substrate peptide (sequence: AEEEIYGEFAKKK) (60). Reactions were initiated through the addition of kinase at a final concentration of 33 nM, and the decrease in absorbance was monitored over 10 min at 30°C in a spectrophotometer (SpectraMax 340PC384). The background activity of the proteins at different drug concentrations was determined in experiments without the substrate peptide and subtracted from the kinase assays with the substrate peptide. To measure the IC_50_ for imatinib, imatinib was serially diluted three-fold in 100% DMSO and added to the reactions to produce an 8-point dose-response curve ranging up to 1 μM. The activity measured by the slope of NADH oxidation (mOD/min) was plotted against the logarithm of the concentration of drug and fit to equation (1) with the hill slope set to 1.

### K_M_ for ATP Kinase activity assay

Abl wt and Abl N368S kinases were phosphorylated overnight. Phosphorylated Abl N368S, Abl wt, and phosphorylated Abl wt kinases were buffer exchanged to 100 mM Tris (pH 8.0), 10 mM MgCl_2_, 1 mM sodium orthovanadate using PD10 gravity columns (Sigma, GE28985255). Using similar kinase activity assay as described above, reactions contained 100nM of kinase protein, 100 mM Tris (pH 8.0), 10 mM MgCl_2_, 1 mM phosphoenolpyruvate, 0.21 mg/ mL NADH, 75 U/mL pyruvate kinase, 105 U/mL lactate dehydrogenase, and 0.5 mM substrate peptide (sequence: AEEEIYGEFAKKK) (60). Reactions were initiated through the addition of ATP at a range of concentrations that was serially diluted three-fold to generate a 8-point ATP titration curve ranging up to 10 mM. The decrease in 340 nM absorbance was monitored over 10 min at 30°C in a spectrophotometer (SpectraMax 340PC384). The activity measured by the slope of NADH oxidation (mOD/min) was plotted against the concentration of ATP and fit to the Michealis-Menten equation to determine V_max_ and K_M_ of ATP.

### Stopped flow kinetics experiments

We monitored inhibitor binding of imatinib and dasatinib by changes in protein fluorescence (38, 41, 61) using a spectrofluorimeteric ligand-binding assay as described previously (61). Protein was mixed in 1:1 ratio constituting a total volume of 200 uL with at least a ten-fold molar excess of drug to obtain a pseudo-first order condition (62). Changes in protein fluorescence were monitored over time using a rapid mixing stopped flow system (Applied Photophysics RX2000). Protein fluorescence decay upon drug binding was fit to a single exponential equation to obtain the observed rate constant (*k*_obs_) (41). The *k*_obs_ was plotted against drug concentration and the association and dissociation rates were calculated based on the relationship: *k*_obs_= *k*_on_[drug]+ *k*_off_. A range of imatinib concentrations was used from 6 μM to 200 μM. The assay was limited to an imatinib concentration of 200 μM due to imatinib solubility.

### Bosutinib kinetics experiments

Abl kinase domain protein (0.2 μM) was pre-incubated with Imatinib (0.67 μM) in 20 mM Tris pH 8, 0.5 mM TCEP. Bosutinib was added to a final concentration of 5 μM, total volume 1 mL. The mixture was immediately dispensed into a black opaque 96-well plate (100 μL per well) and fluorescence emission over time was monitored at 480 nm, with excitation at either 280 nm or 350 nm, in a spectrophotometer (SpectraMax 340PC384). Protein fluorescence change upon imatinib dissociation was fit to a single exponential equation with or without a sloping baseline to obtain the observed rate constant (*k*_obs_) (41).

### Determination of the dissociation constant K_d_ by fluorescence

The intrinsic protein fluorescence of Abl protein decreases upon imatinib binding and therefore can be used as readout for imatinib binding. We titrated imatinib with 18 different concentrations ranging linearly from 0 to 190 nM. Changes in protein fluorescence were measured using a Jobin Yvon Fluoromax-4 (Horiba) spectrofluorimeter: excitation at 290 nm and measuring emission at 340-360 nm at 25°C. The final protein concentration was 50 nM in a buffer containing 20 mM Tris (pH 8.0), 250 mM NaCl, 5% glycerol, 1 mM DTT. Fluorescence versus drug concentration was fit to the following equation: Y=Bmax*X/(K_d_+X) + NS*X + Background, where Bmax is the maximum specific binding, K_d_ is the equilibrium binding constant, NS is the slope, and Background is the amount of nonspecific binding that represents counter background. Triplicates were collected for each protein.

### SAMS molecular dynamics simulations

We selected PDB 2HYY chain A and 2GQG chain A as starting structures for the Abl kinase domain. The N368S mutation was manually introduced using UCSF-Chimera (63). Models were solvated in TIP3P water (64) and neutralized with minimal NaCl. Small molecule ligands were parameterized using the GAFF 1.81 force field (65) and the AM1-BCC charging method (66) from antechamber implemented in the AmberTools software package (67). Molecular dynamics simulations with SAMS enhanced sampling scheme were run using the Amber14SB forcefield (68) through the OpenMM package v7.4.1 (69). Short equilibration (20 ps) was performed before a production run using the OpenMM Langevin integrator in 310.15 K and 1.0 atm with a timestep of 4 fs (using heavy hydrogens assigned a mass of 4 atomic mass units). The implementation of SAMS used here is available on GitHub: https://github.com/choderalab/sams. The SAMS scheme adaptively applies biasing potentials to structural features of interest (dihedrals and distances), and two slightly different strategies were used here. In Strategy 1, we included the seven dihedrals and two distances around the DFG motif sufficient to distinguish discrete activation loop conformations described in Modi and Dunbrack, 2019 (70), and in Strategy 2, we also included two more distances (αC-Glu(+4)-Cα : DFG-Asp-Cγ, and β3-Lys-Cα : DFG-Asp-Cγ) in an attempt to further accelerate transitions between activation loop conformations. Scripts for carrying out the SAMS simulations described here can be found in the following GitHub repository: https://github.com/choderalab/Abl_kinase_N368S.

### Hydrogen bond analysis

For each of the 12 SAMS trajectories generated, all hydrogen bonds (H-bonds) present in each snapshot were identified using the Baker-Hubbard criterion implemented in the software package MDTraj (71) and H-bonds with the most presence (appear in > 50 / 500 snapshots) were kept. By considering all six trajectories for WT and six for N368S, consensus sets of nine H-bonds (N368- ND2 -- D363-O, N368-ND2 -- A380-O, N368-OD1 -- A365-N, N368-OD1 -- R367-NH1, N368- ND2 -- D363-OD1, N368-ND2 -- D363-OD2, N368-ND2 -- D381-OD1, N368-ND2 -- D381- OD2, N368-ND2 -- H361-NE2) and five h-bonds (S368-OG -- D363-O, S368-OG -- A365-N, S368-OG -- A365-O, S368-OG -- A380-O, S368-OG -- R367-NH1) were identified. Statistical uncertainty was estimated as the variance of the corresponding H-bond indicator function corrected by the effective number of uncorrelated configurations in each trajectory using the software package pymbar (72). Scripts for SAMS simulations mentioned above can be found in the following GitHub repository: https://github.com/choderalab/Abl_kinase_N368S.

## Acknowledgements

We acknowledge support for this work by NIH R35 GM119437 (M.A.S.), NIH F30 CA225172 (A.L.), NIH T32 GM008444 (A.L., A.M.R.), NIH P30 CA008748 (J.D.C.), NIH R01 GM121505 (J.D.C.), the Sloan Kettering Institute and Cycle for Survival (J.D.C.). The content is solely the responsibility of the authors and does not necessarily represent the official views of the National Institutes of Health. The authors are grateful to Brooks Bond-Watts and Chris Jeans of the QB3 MacroLab at UC Berkeley for their work on the high-throughput expression of the Abl mutants. S.K. and B.-T. B. are grateful for support by the SGC, a registered charity (no. 1097737) that receives funds from; AbbVie, Bayer AG, Boehringer Ingelheim, the Canada Foundation for Innovation, Eshelman Institute for Innovation, Genentech, Genome Canada through Ontario Genomics Institute [OGI-196], EU/EFPIA/OICR/McGill/KTH/Diamond, Innovative Medicines Initiative 2 Joint Undertaking [EUbOPEN grant 875510], Janssen, Merck KGaA (aka EMD in Canada and US), Pfizer, the São Paulo Research Foundation-FAPESP and Takeda as well as support from the German translational cancer network DKTK and the Frankfurt Cancer Institute (FCI). The authors are grateful to Victoria R. Mingione for her insightful comments during manuscript editing and preparation.

---

**Figure S1.**
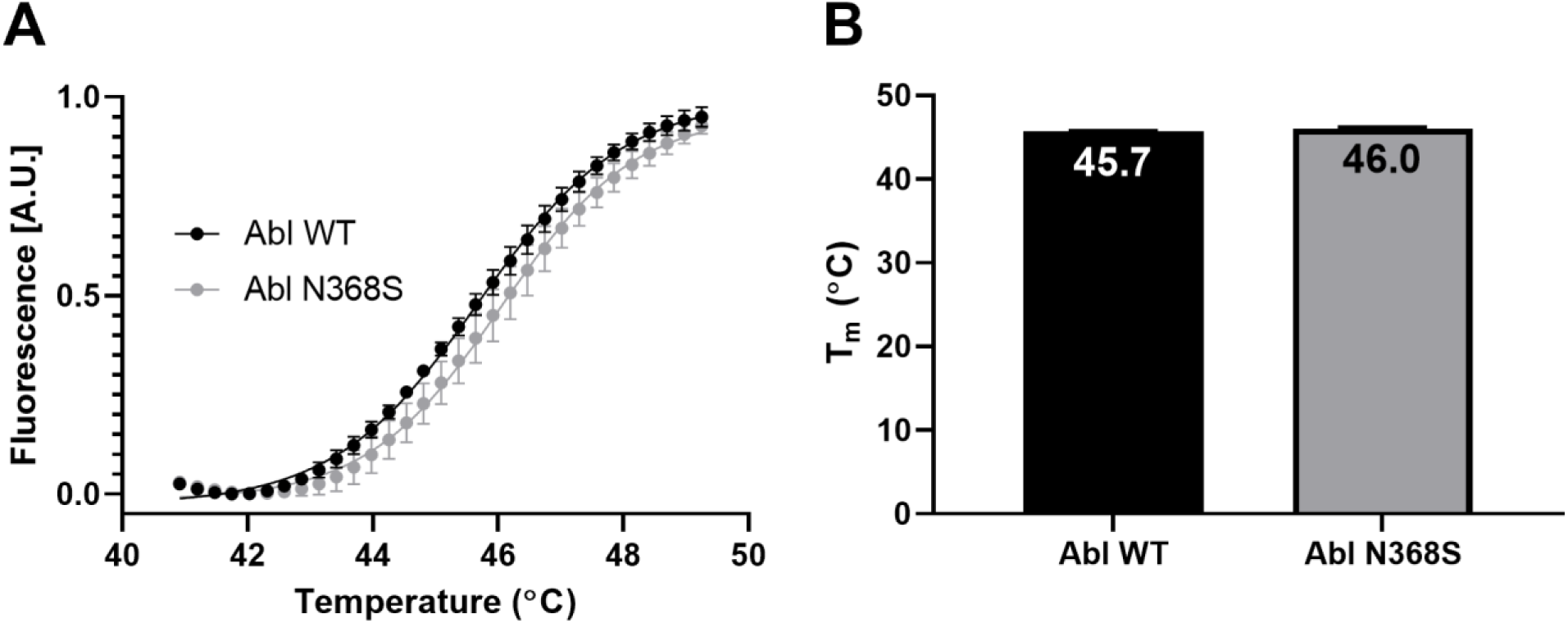
Thermal melt analysis of Abl kinase domain WT and N368S. (A) Representative melt curves of Abl WT and N368S. (B) Thermal melt results. Error bars are the standard deviation of three experiments.

**Figure S2.**
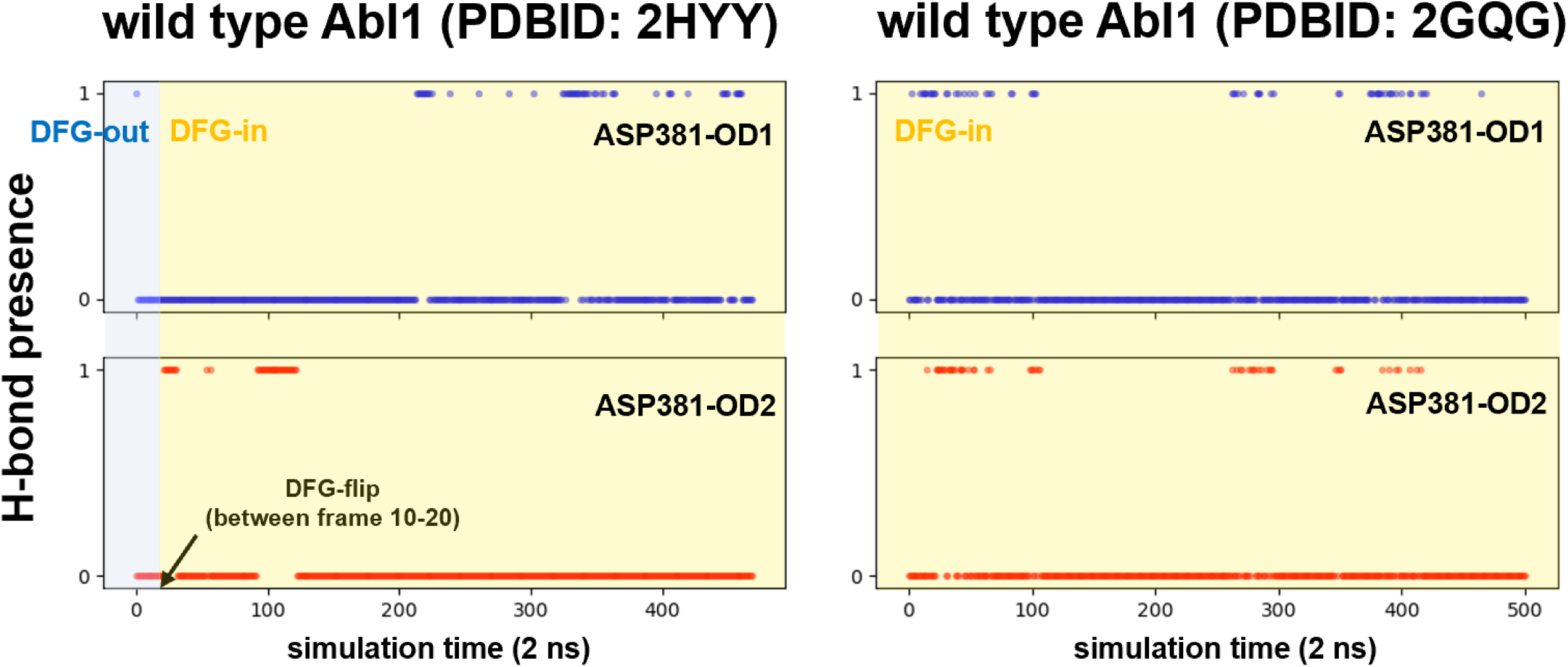
Hydrogen bond presence between N368-ND2 and D381 (either OD1 or OD2) during SAMS simulations. In two wild type Abl1 systems (PDBID: 2HHY and 2GQG) N368 forms H-bond interactions with D381 only in the DFG-in conformation of the kinase (H-bond presence = 1).

**Figure S3.**
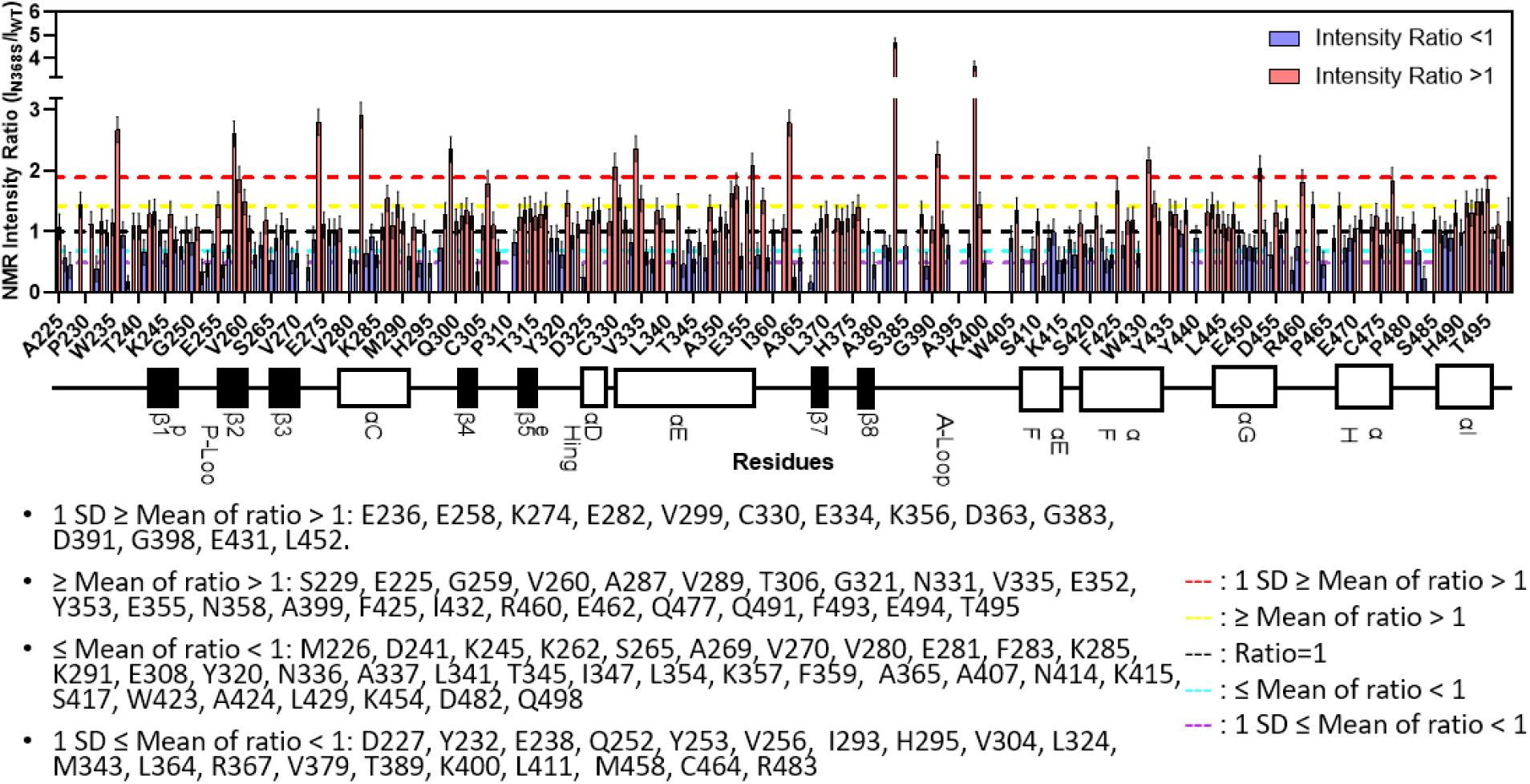
NMR peak intensity ratios. The NMR 1H-15N peak intensity ratios (I_N368S_/I_WT_). The blackline indicates a ratio of 1 (no change of peak intensity upon mutation). The yellow line indicates the mean of all peak intensity values larger than 1 (i.e. peaks are sharper in N368S than in WT). The red line indicates the standard deviation in peak intensity ratios for all values larger than 1. The cyan line is the mean of all peak intensity values small than 1 (i.e. peaks are broader in N368S than in WT). The purple line indicates the standard deviation of all peak intensity rations smaller than 1.

**Figure S4.**
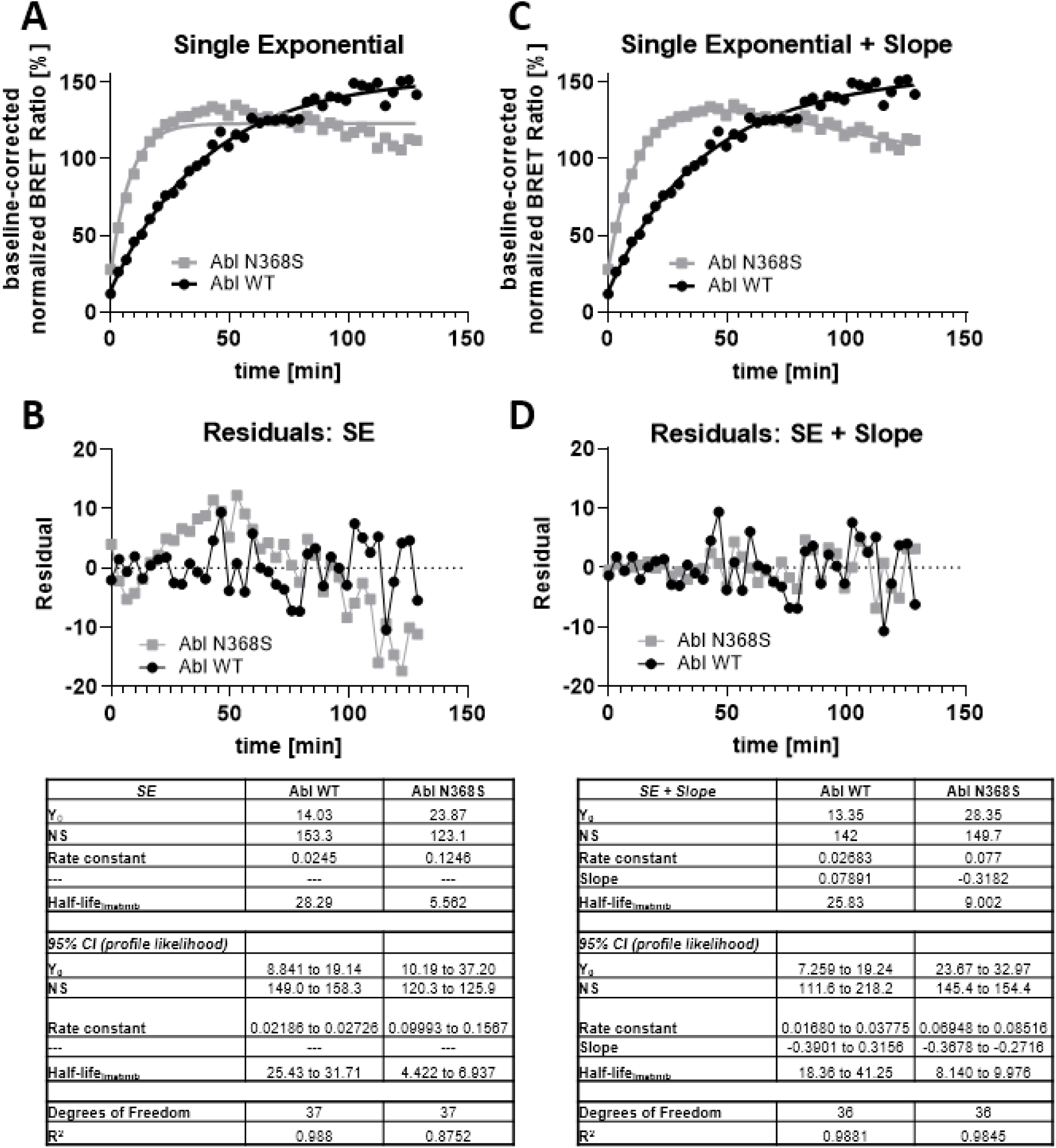
In cell dissociation kinetics of imatinib from Abl WT and N368S. (A) Dissociation kinetics monitored by nanoBRET fit to a single exponential curve. (B) Residuals of single exponential fit to dissociation kinetics and table containing the fitting parameters. (C) Dissociation kinetics monitored by nanoBRET fit to a single exponential curve with a sloping baseline. (D) Residuals of single exponential with sloping baseline fit to dissociation kinetics and table containing the fitting parameters.

